# The lung tissue environment in *Mycobacterium tuberculosis* infection determines local monocyte differentiation

**DOI:** 10.64898/2026.06.25.734601

**Authors:** Alexander Mohapatra, Weihao Zheng, Longhui Qiu, Mark R. Looney, Joel D. Ernst

**Affiliations:** Division of Experimental Medicine, Department of Medicine, University of California, San Francisco, California, USA; Department of Medicine and Department of Laboratory Medicine, University of California, San Francisco, CA, USA

## Abstract

Infection by *Mycobacterium tuberculosis* (Mtb) is characterized by pathogen persistence in lung cells derived from blood monocytes. Since monocyte-derived lung subsets differ in their ability to restrict the growth of intracellular Mtb in mice, understanding the ontogeny of these subsets can inform development of host-directed therapies. Circulating monocytes are proposed to be heterogeneous, arising from distinct bone marrow or spleen progenitors that direct local differentiation. However, the role of the Mtb-infected lung environment in this process has not been addressed. We found that infected and uninfected mice had similar bone marrow monopoiesis, resulting in equivalent monocyte differentiation within the infected lung. While pulmonary Mtb infection also induced splenic monopoiesis, we found no impact on lung monocyte differentiation in splenectomized mice. However, when wildtype monocytes were transferred into Mtb-infected *Sp140^-/-^* recipients, in which excess Type I interferons and neutrophils alter the lung environment, we observed that donor-derived lung subsets resembled recipient-derived cells. In the lungs of Mtb-infected mice, we identified monocyte-derived lung subsets with unique gene expression, associated with specific spatial distributions and cell neighborhoods. These findings suggest that the local lung environment has a larger influence on the phenotypic diversity of monocyte-derived lung cells than does the peripheral environment.

## Introduction

Tuberculosis remains one of the deadliest infectious diseases worldwide, causing more than 1 million deaths annually^1^. Humans who progress to disease following inhalation of *Mycobacterium tuberculosis* (Mtb) exhibit ineffective immune responses, in which infected phagocytes are unable to eliminate the mycobacteria^2^. Since recruited monocyte-derived cells become the major Mtb reservoir in the chronic phase of infection (following T cell recruitment to the lungs)^3–9^, this population is an important cellular target for development of host-directed therapies that can synergize with antimicrobials to achieve faster and more durable Mtb elimination.

We^7,10^ and others^6,9^ have used flow cytometry and single-cell RNA sequencing to define multiple subsets of recruited monocyte-derived cells with differential ability to restrict intracellular Mtb growth. Since these subsets are derived from circulating monocytes produced continuously in the bone marrow (BM)^10^, important questions for therapeutic development are what differentiation pathways underlie the observed diversity in monocyte-derived lung cells and in which anatomical site differentiation occurs. Blood monocytosis is a hallmark of human tuberculosis^11–14^ and is observed in Mtb-infected mice^10^, suggesting that local infection in the lung results in systemic changes in circulating cell populations. Several groups have shown that bone marrow monocytes are heterogeneous in resting mice and are derived from distinct progenitor populations^15–17^. In both mice and humans^14,15,17^, corresponding monocyte subsets have been identified in the blood, and their frequencies can be independently modulated by systemic stimuli based on selective mobilization of monocyte-dendritic cell progenitors (MDPs) or granulocyte-monocyte progenitors (GMPs) in BM.

Therapeutic manipulation of BM monopoiesis to promote “trained immunity” may be a viable strategy to promote enhanced Mtb killing in the lungs. Experimental animal studies in mice^18^ and non-human primates^19^ intravenously inoculated with *M. bovis* bacillus Calmette-Guérin (BCG) suggest that monocytes produced in the setting of BCG replication in BM are functionally altered and possess enhanced ability to restrict Mtb. Conversely, intravenous Mtb infection of BM was shown to impair monocyte function^20^. A separate literature has described “emergency” extramedullary monopoiesis in the spleen in acute myocardial infarction and atherosclerosis^21–23^. Spleen-derived monocytes, as compared to BM-derived cells, in these mouse studies had a distinct contribution to inflammation and wound healing. However, the contribution of BM and spleen monopoiesis to the diversity of recruited monocyte-derived lung cells in pulmonary tuberculosis has not been directly addressed.

To address this question, we characterized monopoiesis in the BM and spleens of mice infected with aerosolized Mtb. We observed an increase in CD319^pos^ monocytes in the BM and blood of Mtb-infected mice. However, BM monocytes from uninfected and Mtb-infected mice yielded similar populations of monocyte-derived lung cells following adoptive transfer into infected recipient mice. While Mtb infection induced splenic monopoiesis, we found no impact on lung monocyte differentiation or Mtb control in splenectomized mice. When transferred into infected recipients with an altered inflammatory lung environment, donor BM monocytes resembled recipient monocytes in their differentiation potential, indicating that local factors influence monocyte differentiation. Within the infected lung, we identified monocyte-derived subsets with unique gene expression, associated with distinct spatial distribution and cell neighborhoods. These findings suggest that the local lung environment has a greater impact on the phenotypic diversity of monocyte-derived lung cells than does the environment of the bone marrow or spleen.

## Results

### Mtb infection alters the blood monocyte population

To determine whether recruited monocyte-derived cell subsets in pulmonary tuberculosis are related to distinct monocyte populations in the circulation, we first assessed the expression of the cell surface proteins CD319 and CD177. These markers were shown to define blood monocyte subsets with distinct BM progenitors (either MDPs or GMPs)^17^. Wildtype mice were infected with aerosolized Mtb, and blood monocytes were analyzed by flow cytometry. In monocytes from uninfected control mice, a CD319^pos^ population was detected, and the frequency of this population was higher in infected mice (52% versus 22% in uninfected controls), accompanied by an increase in the median fluorescence intensity (MFI) of total monocytes (11-fold increase in MFI; Fig. 1a-b, Supplementary Fig. 1a-b). CD177^pos^ monocytes were not detected, although CD177 MFI was higher in blood neutrophils from infected mice.

**Figure 1:**
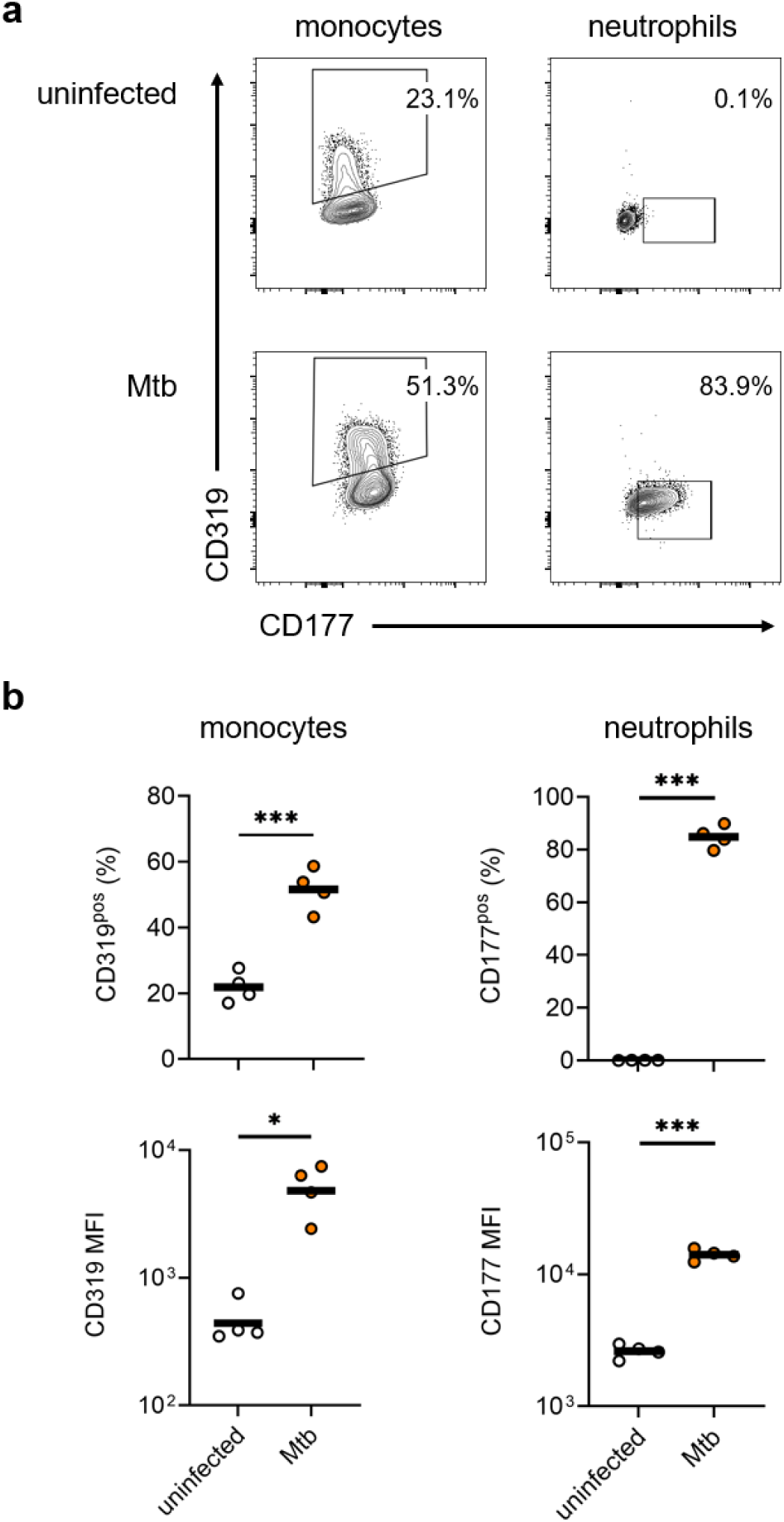
Mtb infection induces expression of CD319 in blood monocytes. Wildtype mice were infected with a target dose of 100 colony-forming units (CFUs) of Mtb, then euthanized at 4 weeks post-infection for analysis of blood cells by flow cytometry (gating in Supplemental Fig. 1). (a) Representative contour plots of the indicated populations from the blood of infected and age-matched, uninfected mice. (b) Frequencies of CD319^pos^ and CD177^pos^ cells (top row) and median fluorescence intensities (MFIs; bottom row) in the indicated populations. Data are from 1 of 2 experiments with similar results, each conducted with 4 mice per condition. Horizontal bar represents the arithmetic mean in all graphs. Significance assessed by Welch’s *t* test. *, *p* <0.05; **, *p* <0.005; ***, *p* <0.0005.

### Bone marrow monocyte and monocyte progenitor numbers are not altered by pulmonary Mtb infection

The observation that CD319^pos^ blood monocytes increase in Mtb-infected mice, and the prior findings that mice^10^ and humans^11–14^ with tuberculosis exhibit blood monocytosis, suggested that pulmonary Mtb infection might alter BM progenitor populations and monocyte production, as was observed in earlier studies of mice with intravenous mycobacterial infection^18,20^. We analyzed BM cells from Mtb-infected mice and uninfected controls and confirmed a higher frequency of CD319^pos^ monocytes in animals infected for 4 weeks (Fig. 2a, Supplementary Fig. 1b). This was associated with BM Mtb burden which was detectable but 10^4^-fold lower, on average, by colony-forming units (CFUs) than that observed in corresponding lung homogenates (Fig. 2b). However, when we used canonical markers to quantitate the number of common myeloid progenitors (CMPs), MDPs, GMPs, or monocytes, no significant differences were observed between uninfected and Mtb-infected mice (Fig. 2c, Supplementary Fig. 2a).

**Figure 2:**
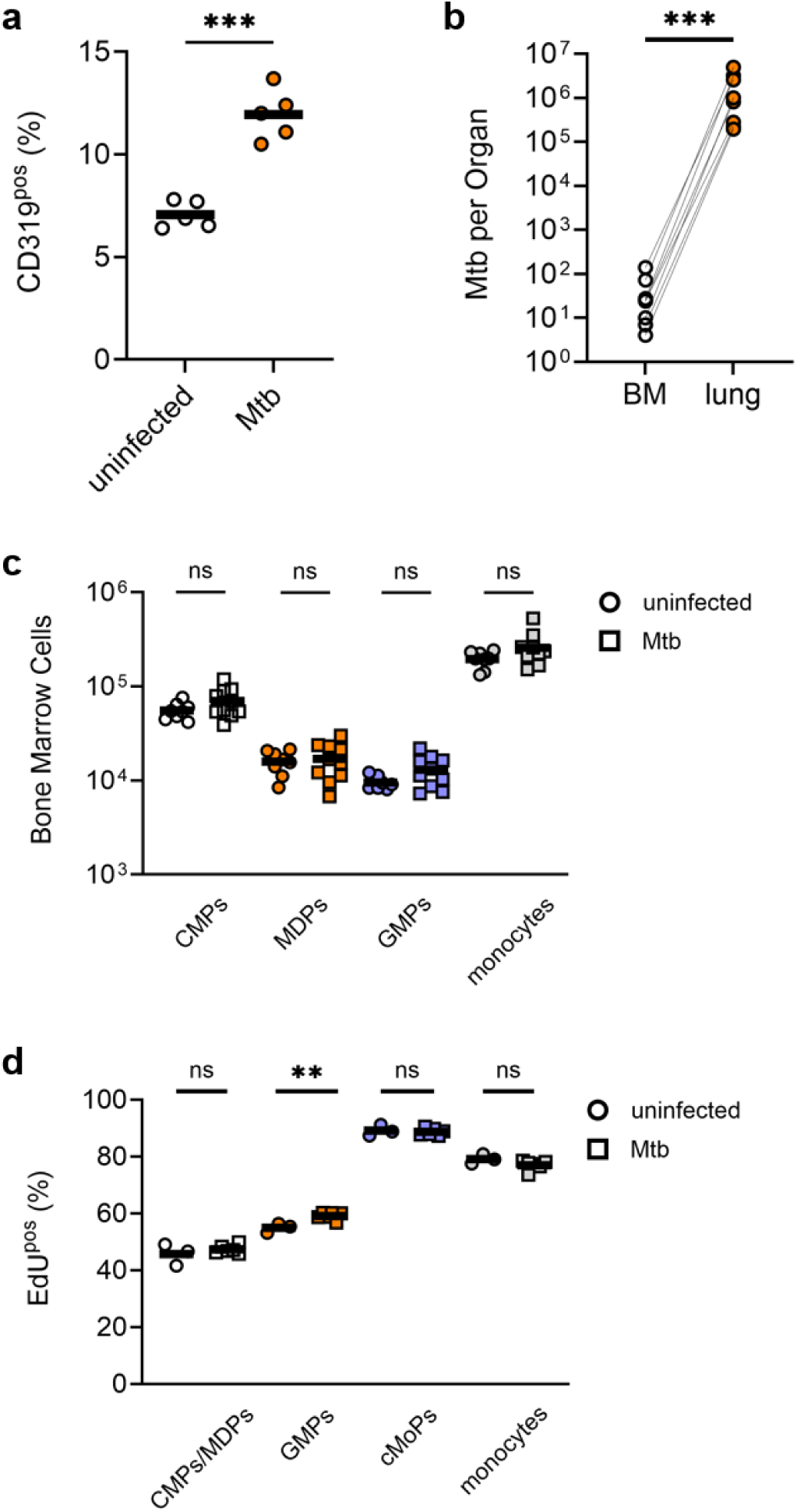
Pulmonary Mtb disseminates to bone marrow but does not alter monocyte or monocyte progenitor numbers. Wildtype mice were infected with ∼100 CFUs of aerosolized Mtb, then euthanized at 4 weeks post-infection for analysis of BM cells by flow cytometry. (a) Frequency of CD319^pos^ cells among total monocytes. (b) Quantitation of Mtb bacilli in the BM and lungs of individual infected mice by colony-forming units (CFUs) on agar. (c) Frequencies of monocytes and progenitors among live BM cells. (d) Infected mice were injected with EdU 48-and 24-hours prior to euthanization. Graph shows frequency of EdU^pos^ cells in the indicated populations. Data are from 1 of 2 experiments with similar results, each conducted with at least 3 mice per condition. Data in (a-c) are compiled from 2 experiments. Horizontal bar represents the arithmetic mean in all graphs. Significance assessed by Welch’s *t* test.

Since this finding could be explained by a combination of increased proliferation of progenitors and increased BM egress^10^, we directly quantitated cycling cells that incorporated 5-ethynyl-2’-deoxyuridine (EdU) injected 48 and 24 hours prior to harvest. We observed a small but significant increase in EdU^pos^ GMPs in Mtb-infected mice, but this was not associated with a corresponding increase in EdU^pos^ BM monocytes (Fig. 2d, Supplementary Fig. 2b). Frequencies of EdU^pos^ cells among other BM progenitor populations were unchanged between uninfected and Mtb-infected mice. Thus, we observed a change in bone marrow monocyte composition, based on CD319 expression, but no change in monocyte production.

### Mtb infection induces splenic monopoiesis but the spleen is not required for Mtb control

We next considered the hypothesis that extramedullary monopoiesis in the spleen contributes to the elevated pool of CD319^pos^ monocytes in the blood of Mtb-infected mice, since the spleen is an established site of Mtb dissemination^24^. We observed significant increases in splenic monocytes and monocyte progenitors in mice infected with Mtb for 4 weeks (Fig. 3a, Supplementary Fig. 1b, 3a), concordant with other studies of inflammation-induced splenic monopoiesis^21–23^. This increase in monocytes occurs after Week 2 of infection (Supplementary Fig. 3b) and is accompanied by splenic lymphocyte proliferation (Supplementary Fig. 3c). To determine whether splenic monopoiesis directly contributes to the accumulation of recruited monocyte-derived cells in the lungs and to Mtb control, we analyzed splenectomized mice infected with Mtb for 4 weeks^25^. By flow cytometry, we identified 3 monocyte-derived lung subsets (Fig. 3b, Supplementary Fig. 1b), termed DN (CD11b^pos^ MHC-II^neg^, CD11c^neg^), MNC1 (CD11b^pos^ MHC-II^pos^, CD11c^neg^), and MNC2 (CD11b^pos^ MHC-II^pos^, CD11c^pos^)^10^. We previously showed that DN cells are rarely infected with Mtb, while MNC1 cells harbor significantly more viable Mtb than MNC2 cells^7^. Mtb-infected, splenectomized mice (spx) had similar numbers of lung MNC1 and MNC2 cells compared to control mice (Fig. 3c). Lung Mtb burdens did not differ in splenectomized mice, when compared to controls (Fig. 3d).

**Figure 3:**
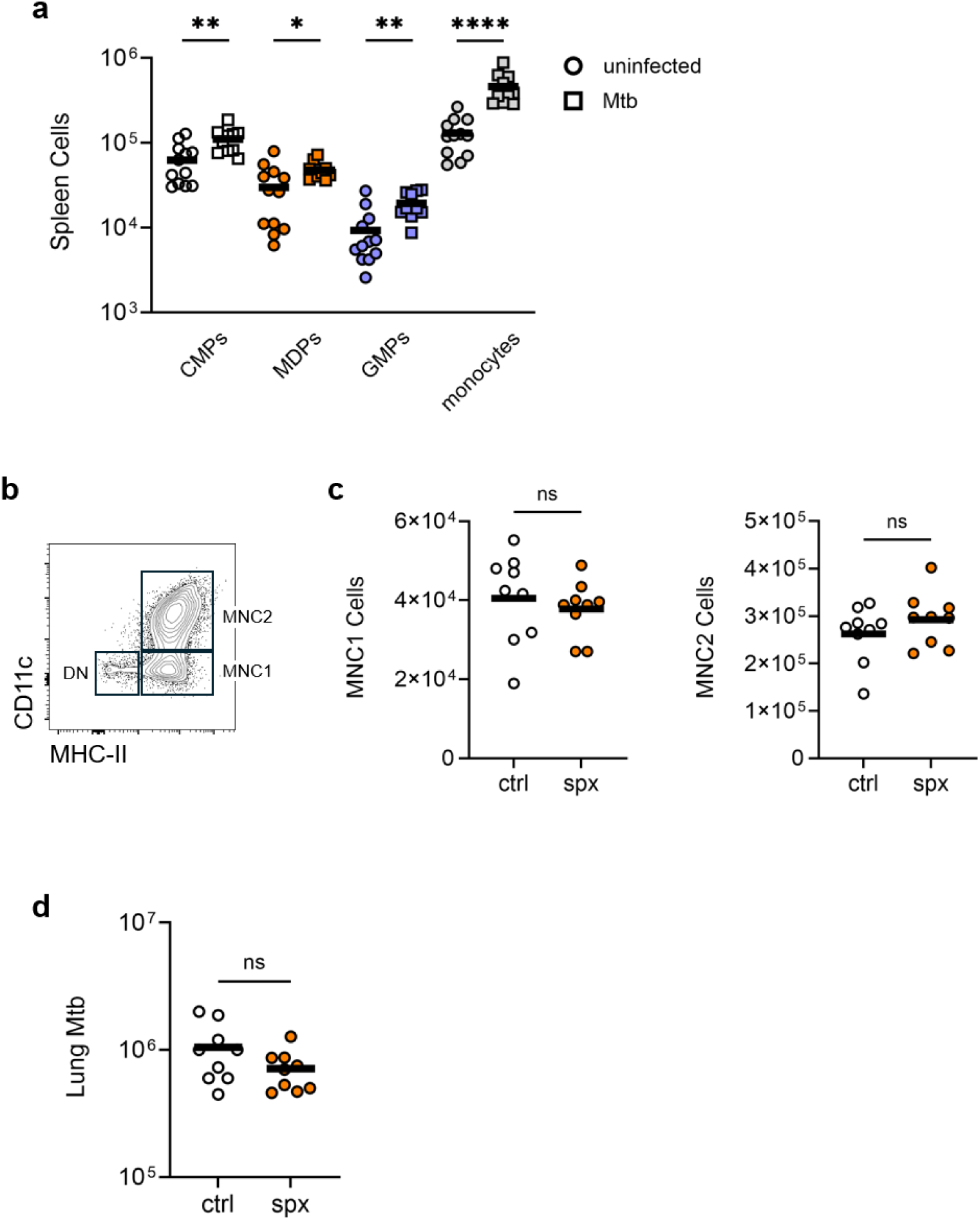
Mtb infection induces splenic monopoiesis but these monocytes are not required for Mtb control. Wildtype mice were infected with ∼100 CFUs of aerosolized Mtb, then euthanized at 4 weeks post-infection for analysis of splenocytes by flow cytometry. (a) Frequencies of monocytes and progenitors among live splenocytes. Data are compiled from 3 experiments. (b) Representative contour plot of monocyte-derived cells in the lungs of Mtb-infected mice pregated to exclude intravascular cells, lymphoid cells, and alveolar macrophages and distinguished by CD11c and MHC-II. (c-d) Mice underwent splenectomy 4 weeks prior to Mtb infection (spx) and were analyzed together with age-matched mice that did not receive splenectomy (ctrl) at 4 weeks post-infection. (c) Quantitation of MNC1 and MNC2 cells in the lungs of infected mice. (d) Quantitation of Mtb bacilli in the lungs. Data are compiled from 2 experiments. Horizontal bar represents the arithmetic mean in all graphs. Significance assessed by Welch’s *t* test.

### Monocytes from uninfected and infected mice differentiate similarly in the lung

Our analyses of bone marrow and spleen monopoiesis in mice with pulmonary Mtb infection failed to explain blood monocytosis but did suggest that the increase in CD319^pos^ blood monocytes with infection alters the population of monocyte-derived cells that are recruited to the lungs and become infected with Mtb. To test the hypothesis that the circulating monocyte pool in infected mice is qualitatively distinct from that of uninfected mice, we isolated BM cells from uninfected and Mtb-infected donor mice congenic at the *Cd45* allele and intravenously transferred equal numbers into congenic recipient mice infected with Mtb for 4 weeks. Three days post-transfer, we analyzed the lungs of recipient mice and identified monocyte-derived cells from uninfected donors, infected donors, and recipients by flow cytometry (Fig. 4a). We confirmed that CD319 expression at this time point remained higher in circulating monocytes from infected donor mice compared to those from uninfected donors (Supplementary Fig. 4a). Monocyte-derived subsets were classified both by CD11c and MHC-II and by CD16.2 and CD206. Subsets classified by these latter surface proteins have distinct gene expression profiles and localization within the lungs^26–28^. Recent work has also suggested that MDPs and GMPs have varying ability to contribute to lung subsets defined by CD16.2 and CD206, depending on the inflammatory context^17^. We observed that donors from uninfected and Mtb-infected mice equivalently differentiated into DN, MNC1, and MNC2 cells in the lungs of Mtb-infected recipients (Fig. 4b). In contrast to infection with influenza virus, in which a substantial CD16.2^neg^ CD206^pos^ recruited monocyte-derived population was observed^17^, we found in pulmonary Mtb infection that all CD206^pos^ cells were also CD16.2^pos^ and classified subsets as DN (CD16.2^neg^ CD206^neg^), CD206^neg^ (CD16.2^pos^ CD206^neg^), or CD206^pos^ (CD16.2^pos^ CD206^pos^) (Fig. 4a). However, we still observed that donor origin did not affect the tendency to differentiate into these subsets (Fig. 4c). Phenotypes of lung cells derived from donor monocytes were similar when BM monocytes purified by magnetic cell separation (MACS) were transferred into infected recipients (Supplementary Fig. 4b-c). Regardless of origin, donor monocytes predominantly differentiated into CD206^neg^ MHC-II^pos^ cells, suggestive of localization within the alveolar interstitium^28^. Collectively, these data suggest that differences in the circulating monocyte pools in uninfected and Mtb-infected mice (Fig. 1b, Supplementary Fig. 4) do not explain the observed frequencies of monocyte-derived lung subsets.

**Figure 4:**
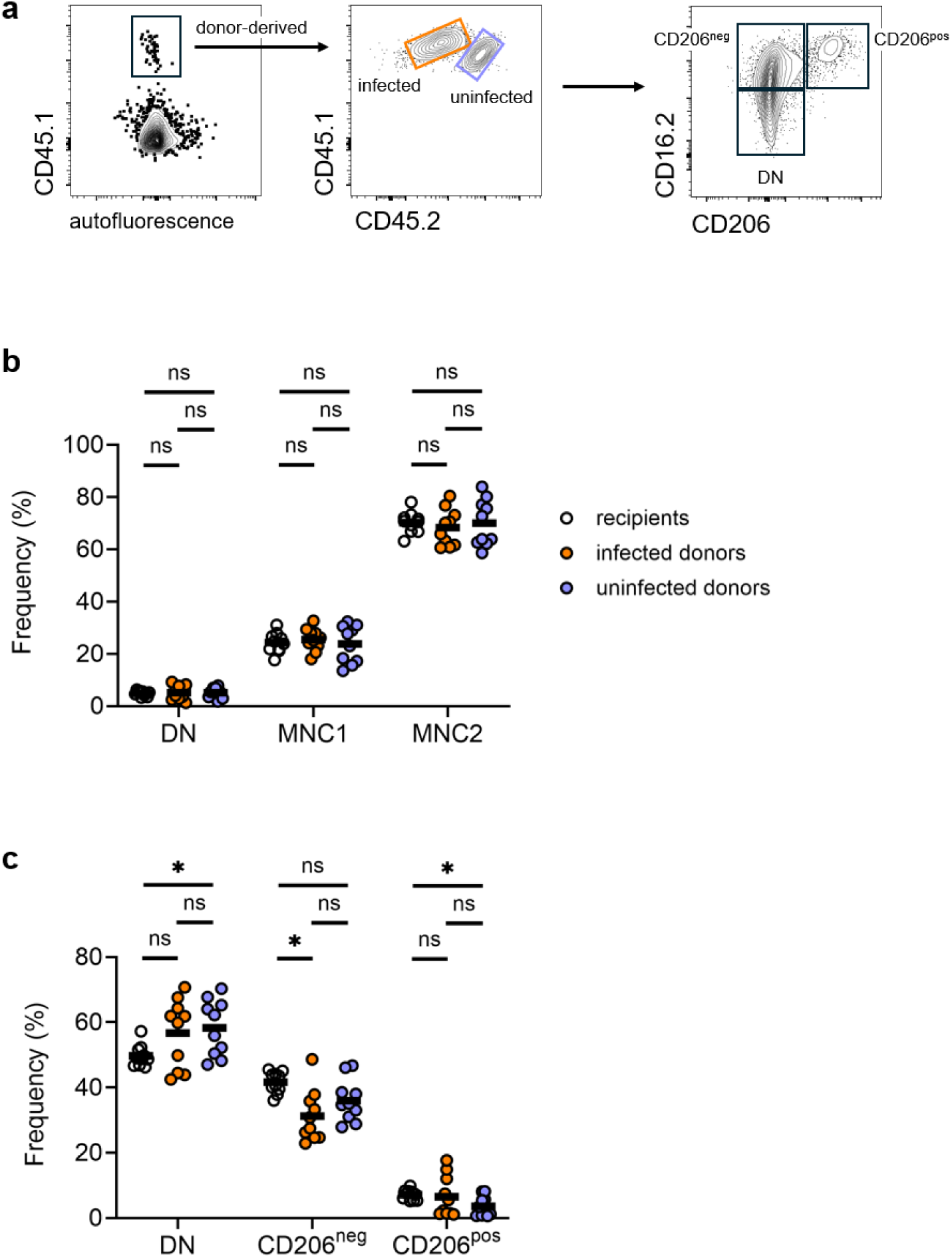
Bone marrow monocytes from uninfected and Mtb-infected mice differentiate into similar lung populations. A *Cd45* congenic strain was infected with ∼100 CFUs of aerosolized Mtb for 4 weeks. Bone marrow cells were isolated from infected mice (CD45.1/CD45.1) and uninfected controls (CD45.1/CD45.2) and intravenously transferred to infected, congenic recipients (CD45.2/CD45.2) also infected for 4 weeks. Lung cells were analyzed by flow cytometry 3 days post-transfer. (a) Representative contour plots of gating to distinguish lung cells derived from uninfected donor cells, infected donor cells, or recipient cells and to characterize these cells by expression of CD16.2 and CD206. (b-c) Frequencies of the indicated populations among monocyte-derived cells, as distinguished by CD11c and MHC-II (b) or CD16.2 and CD206 (c). Data are compiled from 2 experiments. Horizontal bar represents the arithmetic mean in all graphs. Significance assessed by Welch’s ANOVA with multiple comparisons.

### Lung inflammation alters monocyte differentiation

To test the hypothesis that the inflamed lung environment associated with Mtb infection influences lung monocyte differentiation, we turned to the *Sp140^-/-^* mouse model. This genetic background has been shown to confer more rapid development of organized, necrotic granulomas during Mtb infection, due to elevated Type I Interferon production from recruited and resident lung populations and enhanced neutrophil recruitment^29–31^. We found that monocytes have a diminished tendency to differentiate into MNC2 cells in infected *Sp140^-/-^*mice (Fig. 5a). We transferred BM cells from uninfected wildtype mice (*Sp140^+/+^*) into either Mtb-infected wildtype or *Sp140^-/-^* recipients and analyzed monocyte-derived lung subsets 3 days post-transfer. The tendency of donor cells to differentiate into DN, MNC1, or MNC2 cells depended on the recipient genotype (Fig. 5b). We quantitated MFIs for antibodies against cell surface proteins CD11b, CD11c, CD16.2, CD206, Ly6C, and MHC-II. Expression of some proteins, such as CD16.2, varied with recipient genotype, while that of others, such as Ly6C, corresponded with donor genotype (Fig. 5c, Supplementary Fig. 5). We then used principal components analysis to integrate expression of these 6 proteins by flow cytometry to further classify monocyte-derived cells. This analysis supports the observation that wildtype donor monocytes transferred into *Sp140^-/-^* recipients differentiate into lung cells resembling the bulk population in *Sp140^-/-^* mice (Fig. 5d) and indicates that the lung environment strongly influences monocyte differentiation in pulmonary Mtb infection.

**Figure 5:**
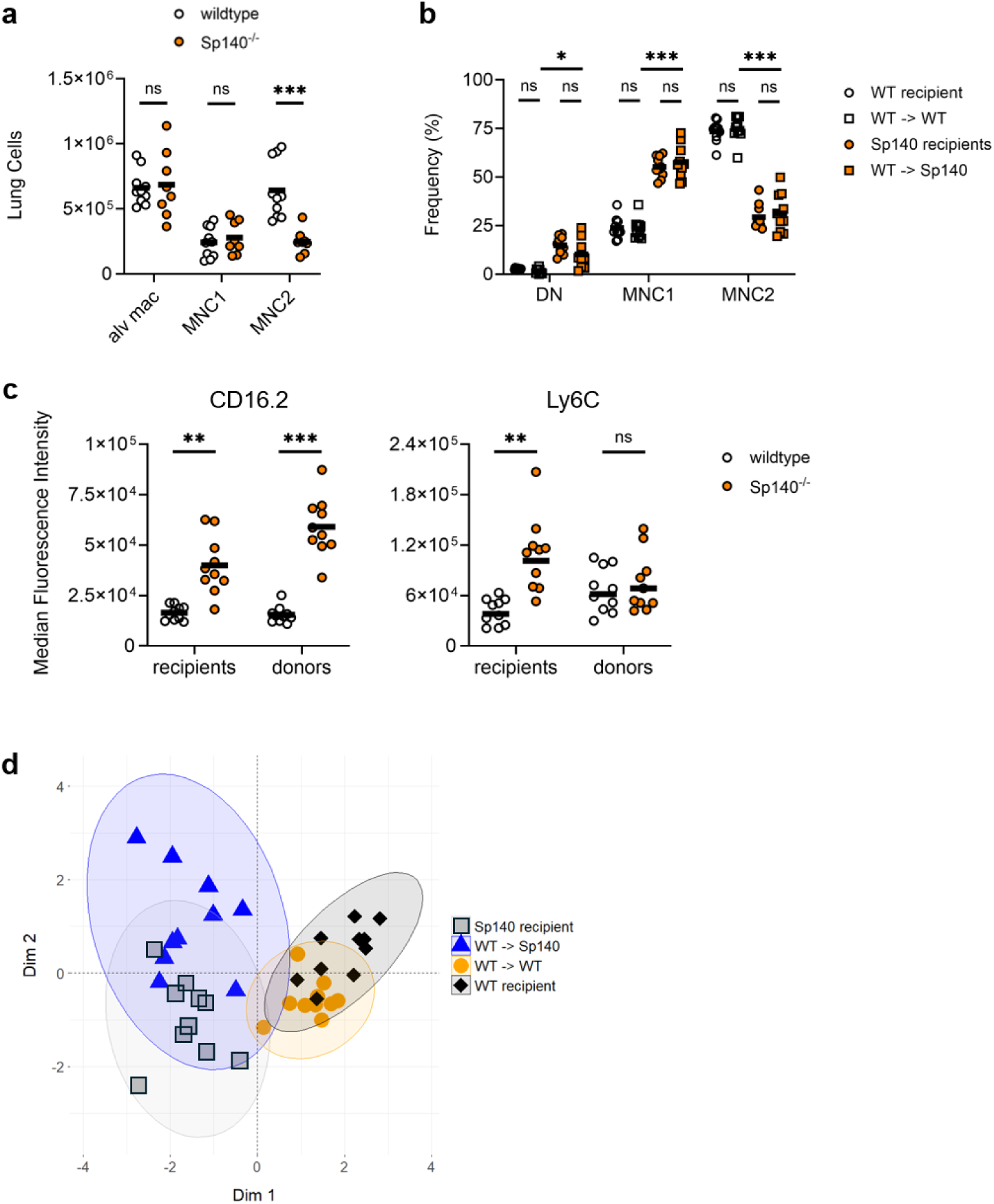
The lung environment has a greater influence than the bone marrow environment on monocyte differentiation in the lungs during Mtb infection. *Cd45* congenic wildtype and Sp140^-/-^mice were infected with ∼50 CFUs of aerosolized Mtb for 4 weeks. (a) Quantitation of lung alveolar macrophages (“alv mac”) and monocyte-derived cells. (b-d) Bone marrow cells were isolated from uninfected wildtype donor mice (CD45.1/CD45.1) and intravenously transferred to congenic recipients (CD45.2/CD45.2) of the indicated genotypes infected for 4 weeks. Lung cells were analyzed by flow cytometry 3 days post-transfer. (b) Frequencies of the indicated populations among monocyte-derived cells. (c) Median fluorescence intensities (MFIs) of CD16.2 and Ly6C on monocyte-derived cells from the indicated bone marrow sources. (d) Principal components analysis of monocyte-derived cells by z-score normalized MFIs of CD11b, CD11c, CD16.2, CD206, MHC-II, and Ly6C. Colors denote bone marrow origin. Ellipses denote 95% confidence interval of clustering. Data are compiled from 2 experiments. Horizontal bar represents the arithmetic mean in all graphs. Significance assessed by Welch’s *t* test or Welch’s ANOVA test with multiple comparisons.

### Monocyte-derived subsets are spatially segregated in the infected lung

Our data indicate a role for the inflamed lung environment in lung monocyte differentiation during Mtb infection. To define gene expression differences between monocyte-derived subsets associated with lung localization, we analyzed lung sections from Mtb-infected mice by spatial transcriptomics. We used several published single-cell RNA sequencing atlases of monocyte-derived lung cells in Mtb-infected mice^8,9,30^ to construct a gene panel interrogated by probes designed for the 10X Xenium platform (Supplementary Fig. 6). We imaged and analyzed spatial transcript expression data for 2 sections each from 2 wildtype mice infected with Mtb for 4 weeks. We used Seurat functions^32^ to define spatial “niches”, regions of an image with unique cell type composition. Using anatomical designations, we defined niches comprising conducting and respiratory bronchi (“airways”), alveolar epithelia (“alveoli”), and the characteristic foci of immune cells surrounding Mtb-infected cells (“foci”) (Fig. 6a, Supplementary Fig. 7a). Ziehl-Neelsen acid-fast staining of slides following Xenium imaging confirmed the presence of Mtb bacilli in immune foci (Fig. 6a). Cell clustering by gene expression and annotation informed by LungMAP^33^ and the reference Mtb atlases identified expected hematopoietic, epithelial, and stromal populations in these lung sections (Fig. 6b, Supplementary Fig. 8) and their prevalence within niches (Fig. 6c).

**Figure 6:**
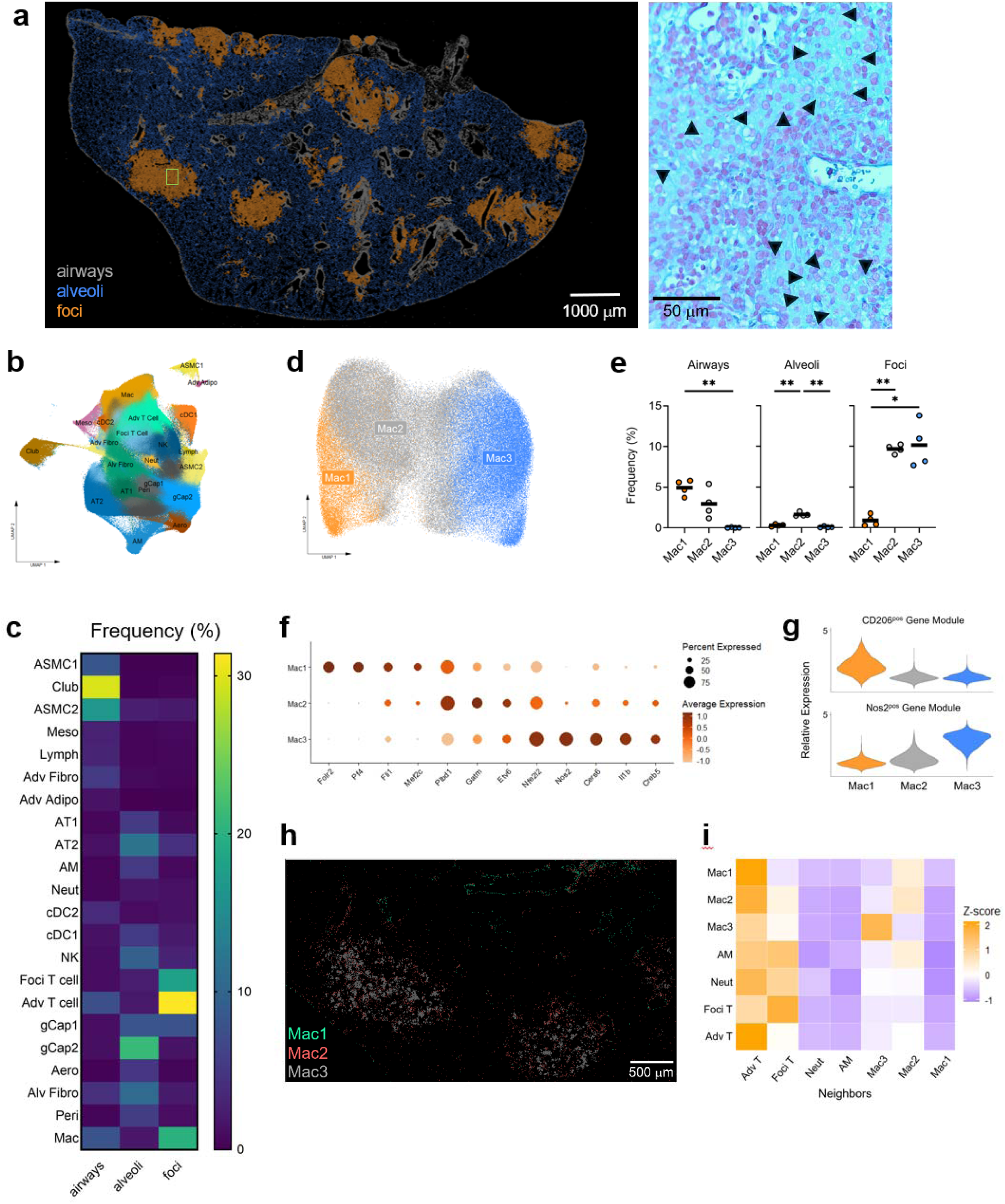
Transcriptionally distinct monocyte-derived cells are spatially segregated in the lungs of Mtb-infected mice. Wildtype mice were infected with ∼100 CFUs of aerosolized Mtb for 4 weeks, then left lung lobes were processed into sections for Xenium spatial transcriptomic analysis. (a) Representative image (left) of niches defined by cells with common gene expression and localization. The region in green is expanded on the right at higher magnification and shows acid-fast staining. Arrow heads denote Mtb bacilli. (b) UMAP showing cell clusters defined by Xenium probe expression (see Methods for description of cell type abbreviations). (c) Heat map of cell cluster frequencies by niche. (d-h) Subclusters of the “Mac” population (macrophages other than alveolar macrophages) were identified. (d) UMAP. (e) Frequencies by niche. (f) Dot plot of select differentially expressed genes. (g) Violin plots of expression of the indicated gene modules. (h) Representative image showing distribution in and around inflammatory foci. (i) Heatmap shows the nearest neighboring cells (columns) to the indicated cell in each row based on the clusters identified in each inflammatory focus, normalized by z-score. Data in (b-g) are compiled from analysis of 2 sections each from 2 infected mice. Data in (a, i) are from 1 section of the 4 analyzed with similar results. Horizontal bar represents the arithmetic mean in all graphs. Significance assessed by Welch’s ANOVA test with multiple comparisons.

We observed that the “Mac” cluster, which includes resident^26–28,34,35^ and recruited monocyte-derived macrophages, but not alveolar macrophages (cluster “AM”), had heterogeneous distribution among niches. Subclustering identified “Mac1”, “Mac2”, and “Mac3” populations with unique gene expression (Fig. 6d) and localization within niches (Fig. 6e). Several genes encoding transcription factors in our Xenium panel exhibited differential expression among Mac subclusters and may relate to lineage specification, such as *Fli1*^36^ and *Mef2c*^36^ in Mac1, *Etv6*^36,37^ and *Nfe2l2*^36^ in Mac2 and Mac3, and *Cers6*^38^ and *Creb5*^36,37^ in Mac3 (Fig. 6f). We also observed differential expression of genes linked to published phenotypes of monocyte-derived “interstitial” macrophages (IMs) in uninfected lungs^26–28,34,35^ and Mtb infection^9,39^. We derived gene signatures from our panel corresponding to these phenotypes and scored signature expression among Mac subsets (Fig. 6g, Supplementary Fig. 9a). These data suggest that Mac1 corresponds to CD206^pos^ Lyve1^pos^ MHC-II^neg^ IMs and are supported by our observation that Mac1 cells tend to associate with airways (Fig. 6e, 6h)^27,28,34^. The predominance of Mac2 and Mac3 cells within immune foci in the alveolar interstitium (Fig. 6e, 6h, Supplementary Fig. 9b) suggests that these cells are similar to reported CD206^neg^ Lyve1^neg^ MHC-II^pos^ IMs^28,35^, though we could not definitively establish this using the genes in our Xenium panel (Supplementary Fig. 9a). Mac3 also corresponds to Nos2^pos^ IMs (Fig. 6g) that have been reported to phagocytose Mtb^9^ and to Mtb-infected macrophages within necrotic granulomas (Supplementary Fig. 9a) in C3HeB/FeJ mice^39^. We also could not conclusively establish whether any of the Mac subclusters was MDP- or GMP-derived based on gene signatures from Trzebanski et al^17^ (Supplementary Fig. 9a).

Within immune foci, we quantified the most common neighbors for each identified cell cluster using SpNeigh^40^ (Fig. 6i, Supplementary Fig. 7b). This analysis demonstrated that Mac1 and Mac2 cells tend to be near a T cell population localized near airways and within immune foci, which we term “adventitial T cells” (Adv T). Conversely, Mac3 cells tend to aggregate within immune foci (Fig. 6h, 6i, Supplementary Fig. 9b). A distinct T cell population found predominantly in immune foci, “Foci T”, is rarely near any of the Mac populations and is more frequently associated with phagocytes such as neutrophils (“Neut”) and AMs, a pattern also observed by Lai et al^6^. Collectively, our observations support the hypothesis that the heterogeneous lung environment of Mtb infection directs monocyte differentiation into subsets with varying ability to engage T cells and control Mtb.

## Discussion

Monocyte-derived lung cells are the dominant Mtb-infected population in the chronic phase of infection (following T cell recruitment to the lungs)^3–7,9^. Since we^7,10^ and others^6,9^ have defined multiple subsets within this population with differential ability to restrict intracellular Mtb growth, we characterized monopoiesis in the BM and spleens of Mtb-infected mice to determine the relative contributions of peripheral versus local differentiation on the phenotypic diversity of monocyte-derived lung cells. We observed an increase in CD319^pos^ monocytes in the BM and blood of Mtb-infected mice. However, BM monocytes from uninfected and Mtb-infected mice yielded similar populations of monocyte-derived lung cells following adoptive transfer into infected recipient mice. We also considered the role of splenic monopoiesis in Mtb infection but found no impact on lung monocyte differentiation or Mtb control in splenectomized mice. When transferred into infected recipients with an altered inflammatory lung environment, donor BM monocytes resembled recipient monocytes in their differentiation potential, indicating that local factors influence monocyte differentiation. Within the infected lung, we identified monocyte-derived subsets with unique gene expression, associated with distinct spatial distribution and cell neighborhoods.

Blood monocytosis is a hallmark of Mtb infection in both humans and mice and is critical to the biology of tuberculosis because Mtb establishes a durable niche in the lung within monocyte-derived cells^3,5–9^. We previously showed that blood monocytosis was associated with increased BM egress in mice infected with aerosolized Mtb^10^. Since BM monocyte numbers were unchanged with infection^10,20^, we hypothesized that increased production would be apparent among monocyte progenitors. However, quantitation of progenitor numbers and proliferation failed to confirm this. Our data suggest that splenic monopoiesis in Mtb-infected mice could additionally contribute to blood monocytosis and are consistent with studies of monocyte recruitment to sites of myocardial infarction in mice^21–23^. Unlike these studies, however, in which spleen-derived monocytes uniquely contributed to myocardial inflammation, we observed that the spleen is a redundant source of monocytes during infection with aerosolized Mtb and not required for lung Mtb control. The observed similarity in lung monocyte differentiation between infected control and splenectomized mice likely explains the unchanged Mtb burdens.

Our data suggest that pulmonary Mtb infection both increases blood monocyte number and alters the composition of this population, based on CD319 expression. Per Trzebanski et al.^17^, this change in expression implies a decrease in the ratio of GMP-derived to MDP-derived monocytes in Mtb-infected mice. However, we observed that BM monocytes from uninfected mice differentiated into lung cells with the same cell surface phenotypes as those derived from the monocytes of infected mice and propose that the lung environment has a greater influence on monocyte differentiation.

One explanation for this discrepancy is that CD319 is induced independent of progenitor origin in circulating monocytes during Mtb infection, raising the possibility that BM monocyte progenitor frequencies are not changed due to infection. While several studies have suggested that monocyte progenitor frequencies can be altered by systemically administered microbial stimuli^15,17^, cytokines^17^, and mycobacteria^18,20^ to foster “trained immunity”, the effects of inflammation outside the bone marrow on monopoiesis are less clear. Trzebanski et al.^17^ demonstrated that intranasal influenza infection in mice could induce preferential differentiation of GMP-derived monocytes in the infected lung. However, Khan et al.^20^ observed no changes in BM MDPs or GMPs with an aerosolized Mtb inoculum that yields relatively modest BM dissemination in our experiments and in other studies^20,41,42^.

Another explanation is that despite an increase in circulating MDP-derived monocytes in Mtb-infected mice, the spatial heterogeneity of inflammation in pulmonary Mtb infection imposes a unique differentiation program on recruited monocytes. Our spatial transcriptomics analysis extends prior observations of heterogeneity among monocyte-derived cells^6–10,30^ by linking these transcriptional profiles, including specific transcription factors, to positioning within the lung and to cell neighborhoods. Recent studies of human lung adenocarcinoma^43^ and Mtb-infected mice^39^ have also linked macrophage subsets to distinct spatial niches. Spatial heterogeneity in the context of influenza infection was not assessed in Trzebanski et al^17^. Therefore, a model unifying our observations in pulmonary Mtb infection with their findings is that the local environment induced by influenza infection reinforces the differentiation potential of GMP-derived monocytes, as is suggested by the reaccumulation of MDP-derived IMs following resolution of inflammation.

The finding that the lung environment alters monocyte differentiation into cells that resemble resident IMs raises the possibility that local proliferation of resident IMs^34^ might contribute to the host response to Mtb infection. While our data do not directly address this possibility, our prior work^10,44,45^ and that of others^46–48^ demonstrate that the increase in IM numbers in the Mtb-infected lung is dependent on CCR2-mediated accumulation of circulating monocytes. This is supported by the observation that influenza infection resulted in recruitment of GMP-derived IMs, supplementing the population of MDP-derived resident IMs^17^. These findings suggest that in inflammatory settings associated with substantial monocyte recruitment, the lung IM population diversifies, despite some similarities in the cell surface phenotypes of recruited and resident IMs.

A limitation of our gene and flow cytometry panels is that they do not conclusively identify monocyte-derived cells of MDP or GMP origin, as has been described in influenza infection^17^. Some of this discrepancy is due to the effect of stimulus-specific inflammation on differentiation of monocyte-derived lung cells; the monocyte-derived subsets we identified in Mtb infection are dissimilar from observed subsets in models of *Cryptococcus* infection^49,50^, toll-like receptor ligand administration^28,34,35^, or influenza infection^17^, highlighting the importance of our data in understanding differentiation in the context of Mtb infection. However, our data do not clarify whether the observed localization of these subsets is a result of differentiation or the driver. Moreover, the focal alveolitis we observe at 4 weeks of infection on a wildtype C57BL/6 background becomes more organized over time^51^, raising the possibility of additional repositioning of monocyte-derived subsets. When comparing the monocyte-derived populations we observe in C57BL/6 mice to those in C3HeB/FeJ^39^ or *Sp140^-/-^*mice^29,30^, which develop faster and more necrotic granulomas, we note common transcriptional and cell surface profiles, suggesting that these mouse models reveal conserved monocyte differentiation within the Mtb-infected lung. We propose that further studies of lung differentiation will reveal pathways that can be modulated to enhance Mtb killing by monocyte-derived cells.

## Methods

### Mice

C57BL/6 mice (strain #: 000664) and congenic CD45.1 mice (strain #: 002014) were obtained from The Jackson Laboratory. *Sp140^-/-^* mice were obtained from the laboratory of Russell Vance. Mice aged 8-12 weeks were used for all experiments, and infected mice were housed in the Animal Biosafety Level 3 facility. All animal protocols used here were approved by the University of California, San Francisco, Animal Care and Use Committees. For injection of 5-ethynyl-2’-deoxyuridine (EdU; Thermo), mice received 0.5 mg in 200 μL PBS intraperitoneally at 48 and 24 hours prior to euthanization. For adoptive cell transfers, cells were resuspended in PBS, filtered through 50 μm strainers, and injected intravenously in 150 μL PBS into the retro-orbital sinus.

### Mycobacterial strains, growth, and aerosol infection

*M. tuberculosis* strain H37Rv was grown in Middlebrook 7H9 medium (BD) supplemented with 10% (vol/vol) ADC (albumin, dextrose, and catalase), 0.05% Tween 80, and 0.2% glycerol. Mice were infected with H37Rv via aerosol using an inhalation exposure unit from Glas-Col, as previously described^7,10^. Mid-log cultures were centrifuged at 800 × *g* to pellet clumps. Clump-free cultures were then diluted, and 5 mL inoculum was added to the nebulizer. The target dose was 100 CFUs/mouse. Actual infection dose was determined by plating lung homogenates 24 hours post-infection on 7H11 agar plates and enumerating CFUs after 3 weeks of incubation at 37°C.

### Adoptive cell transfers

For transfer of unfractionated BM cells, femurs and tibias were collected from donor mice and flushed with FACS buffer to extract marrow. After mashing through 70 μm strainers, BM cells were dissolved in ammonium-chloride-potassium (ACK) lysis buffer (Gibco) for red blood cell lysis. Single-cell suspensions were washed and resuspended in PBS with 3% HI-FBS (vol/vol) and 2 mM EDTA (FACS buffer). Live cells were quantitated by Trypan Blue exclusion using the Countess 3 (Thermo). For mixed transfers, cells of each condition were combined in equal numbers and resuspended in PBS at 20 x 10^6^ cells/150 μL for intravenous injection of recipients (see above).

For transfer of BM monocytes purified by magnetic cell separation (MACS; Miltenyi Biotec), BM single-cell suspensions were prepared and quantitated, as above. Cells were processed using the Miltenyi Monocyte Isolation Kit for depletion of non-monocytes, according to the manufacturer’s guidelines. For additional removal of monocyte progenitors, magnetic column flow-through was quantitated and then incubated with biotinylated antibodies to Sca-11 (D7), c-Kit (2B8), and FLT3 (A2F10) for 15 min on ice. Stained cells were then incubated with Miltenyi Anti-Biotin MicroBeads and loaded onto a second magnetic column for depletion, according to the manufacturer’s guidelines. Flow-through was quantitated, and cells were resuspended in PBS at 1.5 x 10^6^ cells/150 μL for intravenous injection.

### Splenectomy

Splenectomy was performed as described in Valet et al^25^. Briefly, an anesthetized mouse was shaved and prepped on the left side with povidone-iodine antiseptic. A 2-cm horizontal skin incision midway between the last rib and the hip joint was made with surgical scissors, followed by a 1-cm incision in the peritoneal wall. The spleen was then gently pulled out with forceps. The splenic artery and vein were ligated with a 7-0 silk suture, and the spleen was removed. The peritoneal wall was closed with 1 absorbable 5-0 silk suture, and the skin was closed with two 5-0 silk sutures. Mice received buprenorphine every 12 hours for 2 days after the operation. Following 4 weeks of rest, splenectomized mice were infected with Mtb, as above, along with an age-matched cohort that did not undergo surgery.

### Tissue harvests and processing for flow cytometry

Mice were anesthetized by inhalation of isoflurane and then retro-orbitally injected with 1 μg of anti-CD45 antibody (clone 30-F11) in 150 μL of phosphate-buffered saline (PBS). Three minutes after antibody injection, mice were euthanized by CO_2_ inhalation and cervical dislocation. Blood was collected from the heart and added to an equal volume of acid citrate-dextrose solution (Sigma). Lungs were collected and placed in Hanks’ balanced salt solution (HBSS) containing 50 μg/mL Liberase (Sigma) and 30 μg/mL DNase I (Sigma). Spleens were collected and placed in FACS buffer. Lungs were incubated at 37°C for 30 min, then processed with a gentleMACS dissociator (Miltenyi) and mashed through 70 μm strainers. Spleens were also mashed through 70 μm strainers. Aliquots for plating on 7H11 agar for CFU enumeration were taken prior to centrifugation. Following centrifugation, lung or spleen cell suspensions were dissolved in ACK buffer for red blood cell lysis. Blood cells underwent red blood cell lysis with ACK buffer twice. Bone marrow suspensions were processed, as described above for adoptive cell transfers. Single-cell suspensions were washed and resuspended in FACS buffer. For staining of surface antigens, cells were washed with PBS, then stained with viability dye Live-or-Dye 750/777 (Biotium) and anti-CD16/32 (clone 2.4G2; Fc receptor blockade) for 15 min at 4°C. After washing, cells were resuspended in FACS buffer with a cocktail of fluorochrome-conjugated antibodies against cell surface proteins, including B220 (RA3-6B2), c-Kit (2B8), CD11b (M1/70), CD11c (N418), CD16.2 (9E9), CD19 (6D5), CD34 (SA376A4), CD90.2 (30-H12), CD127 (A7R34), CD177 (Y127), CD206 (C068C2), CD319 (4G2), CSF1R (AFS98), FLT3 (A2F10), Ly6C (HK1.4), Ly6G (1A8), MHC-II I-A and I-E (M5/114.15.2), NK1.1 (S17016D), Sca-1 (D7), Siglec F (S17007L), and Ter119, for 25 min at 4°C. See Supplementary Figures for gating strategies. Cells were then washed with PBS, fixed with 4% paraformaldehyde (vol/vol) for 30 min at room temperature, and washed again in PBS. Samples were acquired using an Aurora spectral flow cytometer (Cytek Biosciences). Flow cytometry data were analyzed using SpectroFlo 3.3 (Cytek) and FlowJo 10.10.0. GraphPad Prism 10.4.1, FactoMineR 2.12^52^, and factoextra 2.0.0^53^ were used for graphical presentation and statistical analysis.

For detection of intracellular EdU, cells were fixed, permeabilized, and incubated with Thermo Click-iT Plus reagents following staining for cell surface proteins, according to the manufacturer’s guidelines. Phycoerythrin and some tandem fluorochromes were avoided due to incompatibility with Click-iT chemistry.

### Xenium spatial transcriptomics

Mice infected with Mtb for 27 days were euthanized, as above, and 10% neutral buffered formalin was instilled intratracheally to inflate lung lobes. Left lobes were excised and fixed in 5 mL 10% formalin for 24 hours at room temperature. Lobes were transferred out of BSL3 facility and placed in 10 mL 70% ethanol for 24 hours. Following embedding in paraffin blocks, 5 μm sections were cut onto Xenium slides, according to 10X Genomics guidelines. Three sections from each block were placed on a single Xenium slide. Two Xenium slides underwent deparaffinization, decrosslinking, probe hybridization, ligation, amplification, cell segmentation staining, autofluorescence quenching, and nuclei staining according to 10X guidelines. Probes utilized targeted a total of 429 unique genes, including a set specific for a custom 50-gene add-on panel designed by 10X and those contained in the 10X Mouse Tissue Atlassing Panel. The 50-gene custom panel was designed using spapros 0.1.6^54^, using macrophage clusters from reference cells atlases^8,9,30^ processed by scanpy 1.11.5^55^. Slides were imaged on the Xenium Analyzer with software version 4.0.1.4, according to 10X guidelines. Following Xenium Analyzer imaging, the autofluorescence quencher was removed from Xenium slides using aqueous sodium hydrosulfite, according to 10X guidelines. Ziehl-Neelsen acid-fast staining^56^ was then performed to identify Mtb bacilli on Xenium slides. Images were captured using the 40X objective of a Nikon Ti inverted microscope with a DS-Ri2 camera.

### Spatial transcriptomics analysis

Xenium raw data was processed using 10X Space Ranger version 4.0.1. Processed data from 4 sections on 2 Xenium slides were integrated into a single Seurat v5 object, using IntegrateLayers with FastMNNIntegration (Seurat version 5.4)^32^, containing more than 1.3 million cells. Dimensional reduction and clustering were then performed on a subsample of 100,000 cells using SketchData. Clusters were manually annotated using reference cell atlases^8,9,30,33^ and labeled as: adventitial adipocytes (“Adv Adipo”), adventitial fibroblasts (“Adv Fibro”), adventitial T cells (“Adv T cell”), alveolar capillary aerocytes (“Aero”), alveolar fibroblasts (“Alv Fibro”), alveolar macrophages (“AM”), type I (“AT1”) and type II (“AT2”) alveolar epithelial cells, airway smooth muscle cell subset 1 (“ASMC1”) and subset 2 (“ASMC2”), type 1 (“cDC1”) and type 2 (“cDC2”) conventional dendritic cells, club cells (“Club”), immune foci T cells (“Foci T cell”), alveolar capillary cell subset 1 (“gCap1”) and subset 2 (“gCap2”), lymphatic endothelial cells (“Lymph”), monocyte-derived macrophages (“Mac”), mesothelial cells (“Meso”), neutrophils (“Neut”), natural killer cells (“NK”), pericytes (“Peri”). The ProjectData function was then used to apply cluster labels and dimensional reductions to the full dataset. Mac cells from this object were subclustered for further analysis. Expression of gene signatures among Mac subclusters was assessed using AddModuleScore, with 15 control genes per signature gene queried.

For analysis incorporating spatial coordinates, the integrated Seurat object was split by image, and the BuildNicheAssay function was used to identify niches. Subsets of each image object containing only foci niches and select cell types were analyzed using SpNeigh (version 0.99.42) computeSpatialInteractionMatrix^40^. This function quantitates the nearest neighbors (*k* = 10) of each cell in the object. Niche and cluster labels were imported into 10X Xenium Explorer 4.1.1 to generate figure images. Other figure plots were generated using Seurat and scCustomize 3.2.4 (RRID: SCR_024675)^57^ functions.

### Data Availability

The spatial transcriptomics data supporting the findings of this study are available in the Gene Expression Omnibus (GEO) under the accession number GSE336209. Secondary data sources used in this study are available at GSE216023^30^, GSE245950^9^, and GSE263891^8^.

## Acknowledgements

We thank Lucas Chen, Kelley Martinez, and Michael Kwon for technical assistance. The UCSF Division of Experimental Medicine Core Immunology Lab and Parnassus Flow CoLab (RRID: SCR_018206) facilitated the generation of flow cytometry data. We acknowledge the UCSF Center for Advanced Light Microscopy for assistance with image acquisition. We also acknowledge the UCSF Mouse Microsurgery Core for experiments with splenectomized mice.

## Author Contributions

Conceptualization: A.M. and J.D.E.; methodology: A.M., L.Q., M.R.L., and J.D.E.; investigation: A.M., W.Z., and L.Q.; writing – original draft: A.M. and J.D.E.; writing – review and editing: A.M. and J.D.E.; funding acquisition: A.M., W.Z., M.R.L., and J.D.E.; supervision: J.D.E.

