## Supplemental Figures for "The lung tissue environment in *Mycobacterium tuberculosis* infection determines local monocyte differentiation"

Supplementary Figure 1

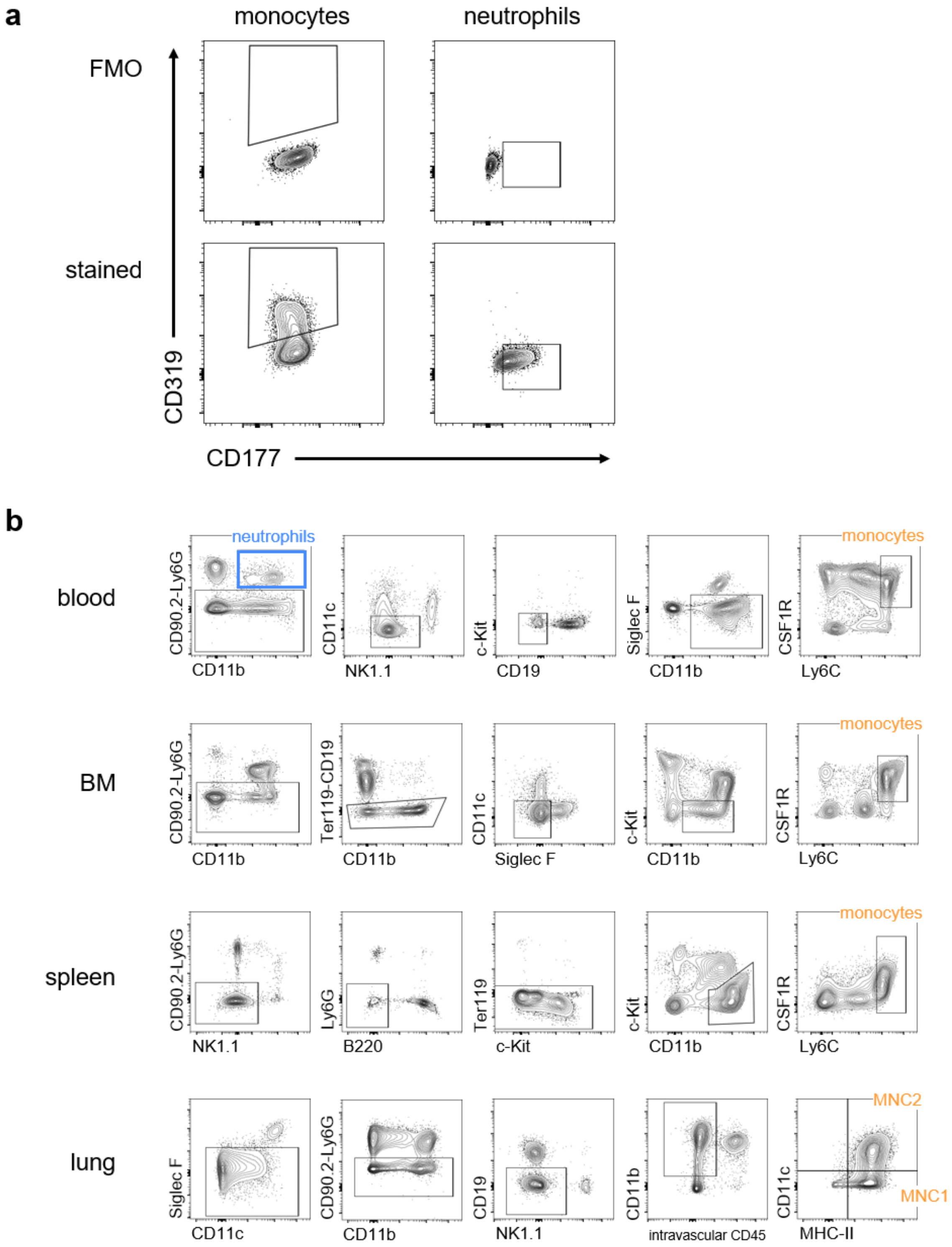


Supplementary Figure 1: Representative flow cytometry of blood, BM, spleen, and lung in Mtb-infected mice. (a) Staining for CD319 and CD177 in blood cells with fluorescence-minus-one (FMO) controls. (b) Gating of monocytes or monocyte-derived cells in the indicated organs.

Supplementary Figure 2

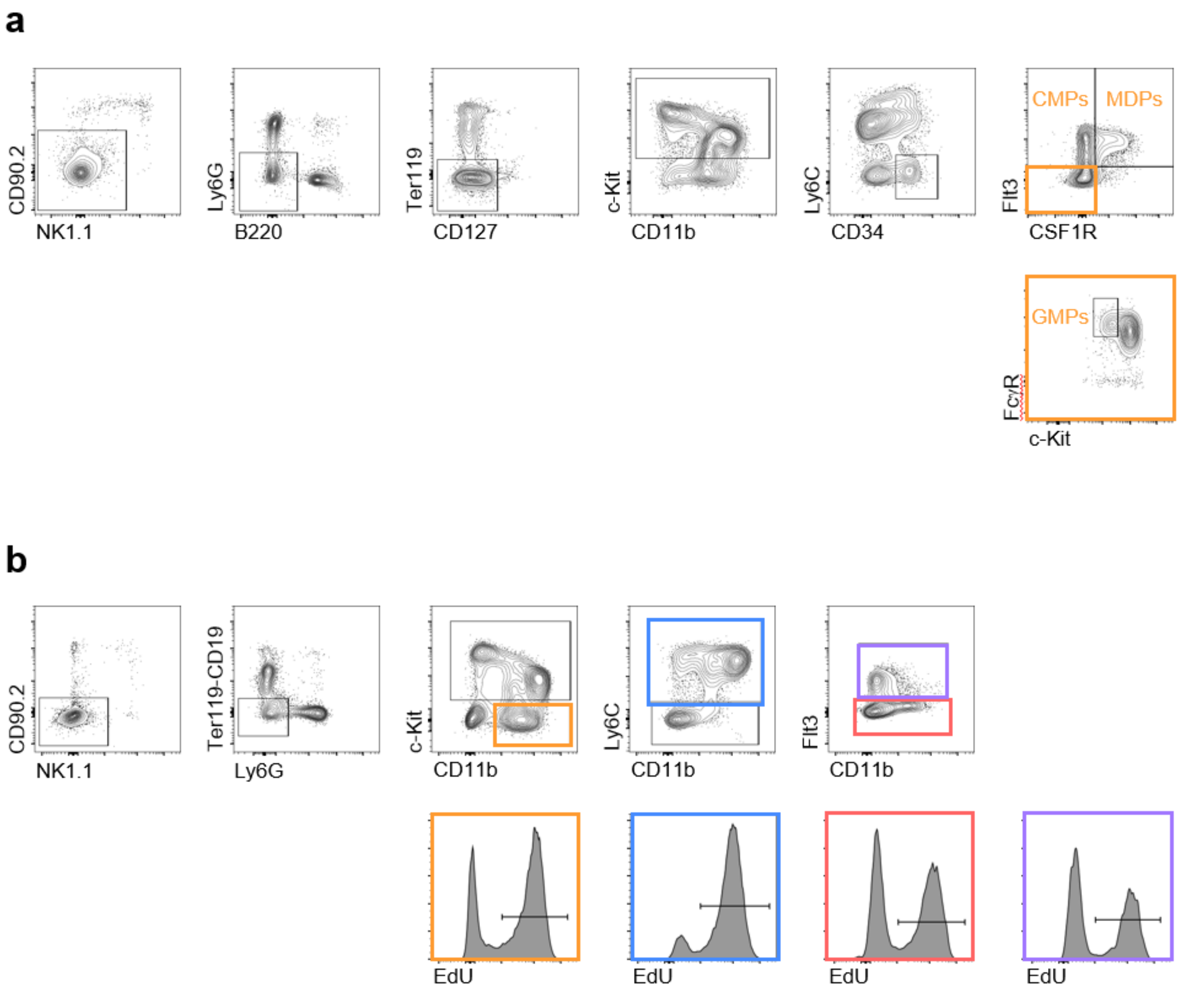


Supplementary Figure 2: Representative flow cytometry of monocyte progenitors in BM of Mtb-infected mice. (a) Gating of progenitors to determine frequency. (b) Gating of progenitors to quantitate EdU-positive cells.

#### Supplementary Figure 3

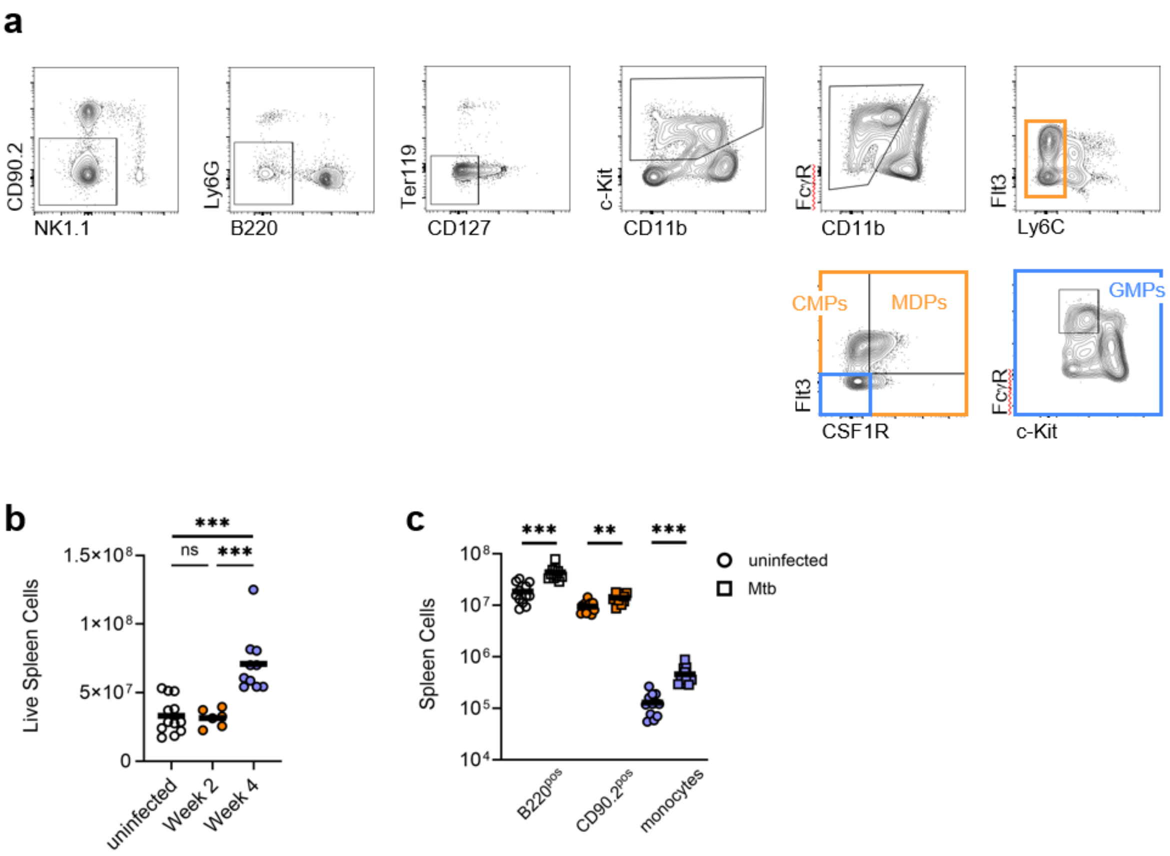


Supplementary Figure 3: Flow cytometry of splenocytes in Mtb-infected mice. (a) Representative gating of monocyte progenitors in the spleen. (b) Quantitation of total live cells in mice infected for the indicated times. (c) Quantitation of the indicated populations in uninfected mice or after 4 weeks of infection. Data in (b-c) are compiled from 2 experiments. Horizontal bar represents the arithmetic mean in all graphs. Significance assessed by Welch's ANOVA with multiple comparisons (b) or Welch's *t* test (c).

Supplementary Figure 4

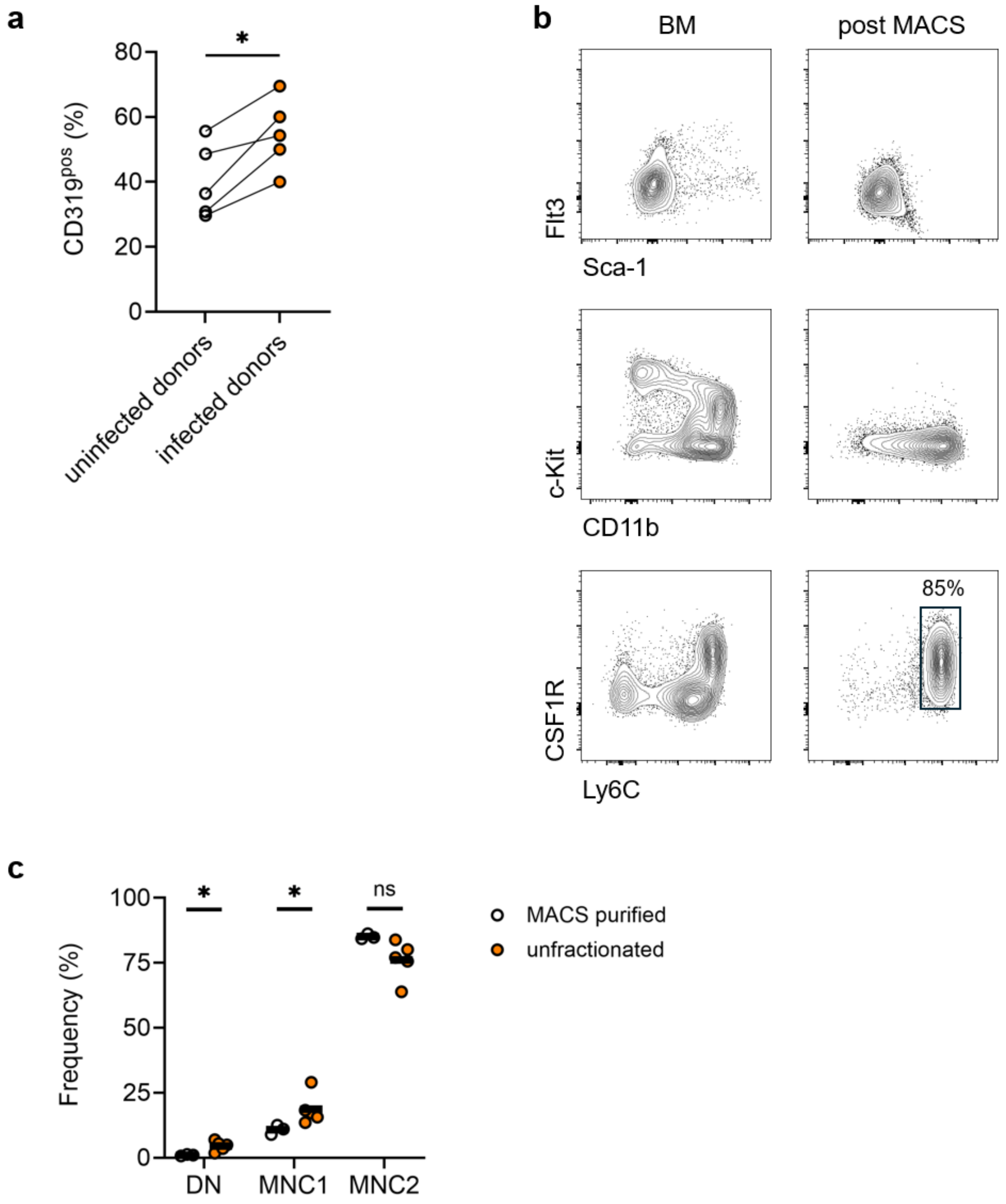


Supplementary Figure 4: Flow cytometry of monocytes transferred into Mtb-infected recipients. (a) BM cells from uninfected mice and mice infected for 4 weeks were co-transferred into infected recipients for 3 days. Expression of CD319 in donor blood monocytes in the blood was higher among monocytes from Mtb-infected mice than in cells from uninfected mice. Data are compiled from 2 experiments. (b) Flow cytometry of BM monocytes purified by magnetic cell separation (MACS). (c) Frequencies of the indicated populations among monocyte-derived lung cells in Mtb-infected recipients 1 week after receiving either MACS purified monocytes or unfractionated BM cells. Data are compiled from 2 experiments. Horizontal bar represents the arithmetic mean in all graphs. Significance assessed by Welch's *t* test for paired samples (a) or unpaired samples (c).

#### Supplementary Figure 5

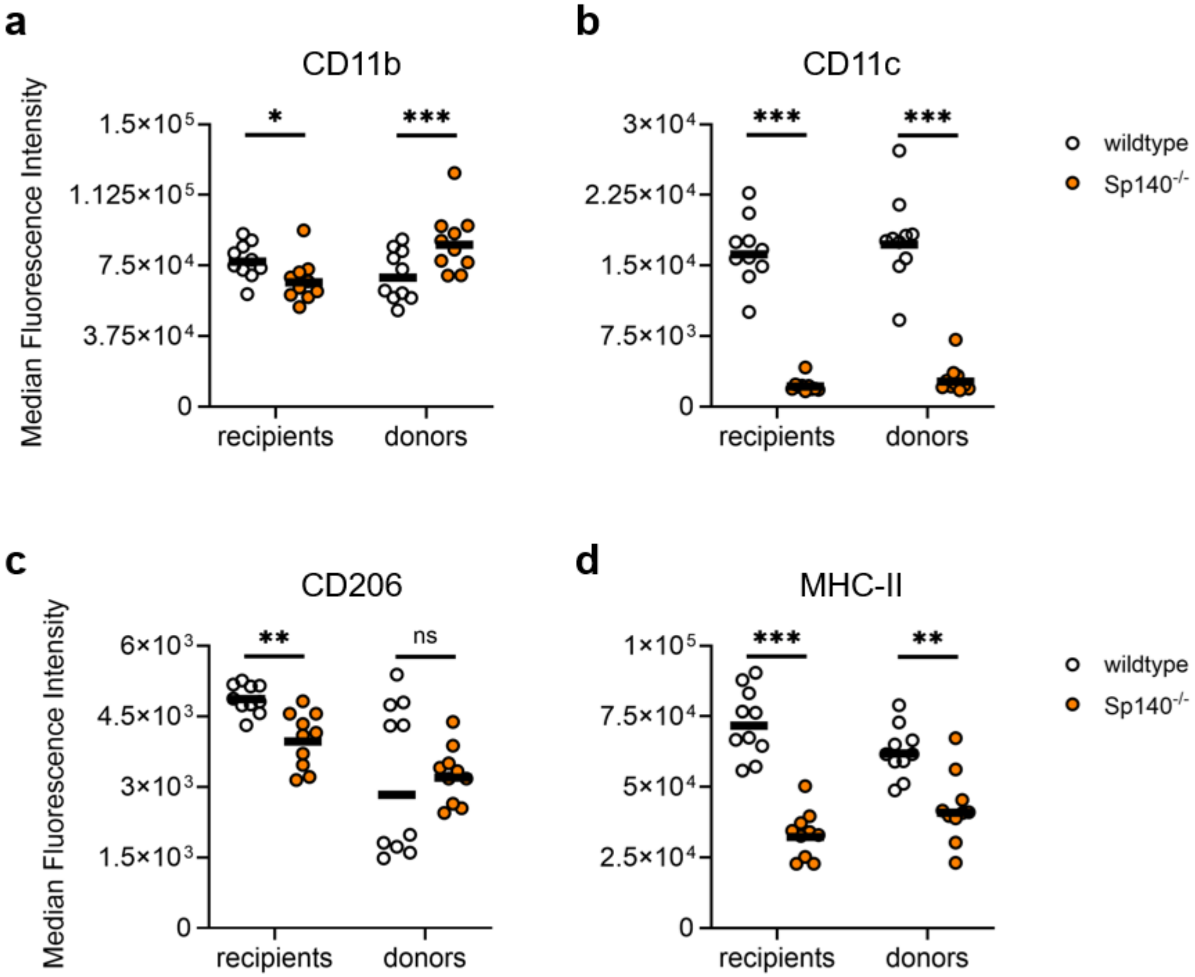


Supplementary Figure 5: Flow cytometry of donor monocyte-derived lung cells in Mtb-infected *Sp140$^{-/-}$* recipients. Median fluorescence intensities of antibody staining for CD11b (a), CD11c (b), CD206 (c), and MHC-II (d). Data are compiled from 2 experiments. Horizontal bar represents the arithmetic mean in all graphs. Significance assessed by Welch's *t* test.

### Supplementary Figure 6

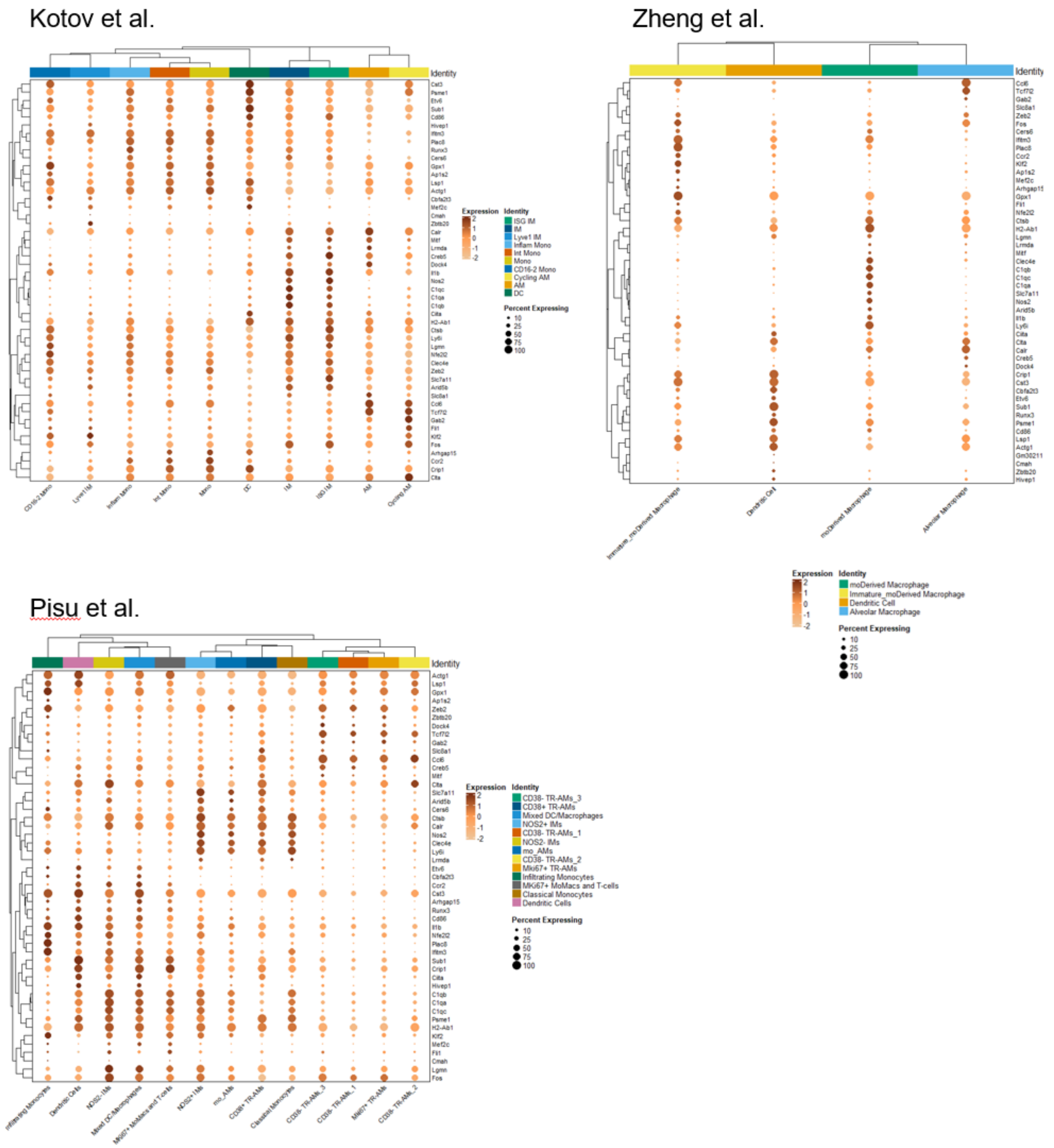


Supplementary Figure 6: Expression of genes in the Xenium add-on custom panel in reference single-cell atlases. Clustered dot plots show relative gene expression and frequency of expressing cells among myeloid populations from the lungs of Mtb-infected mice.

Supplementary Figure 7

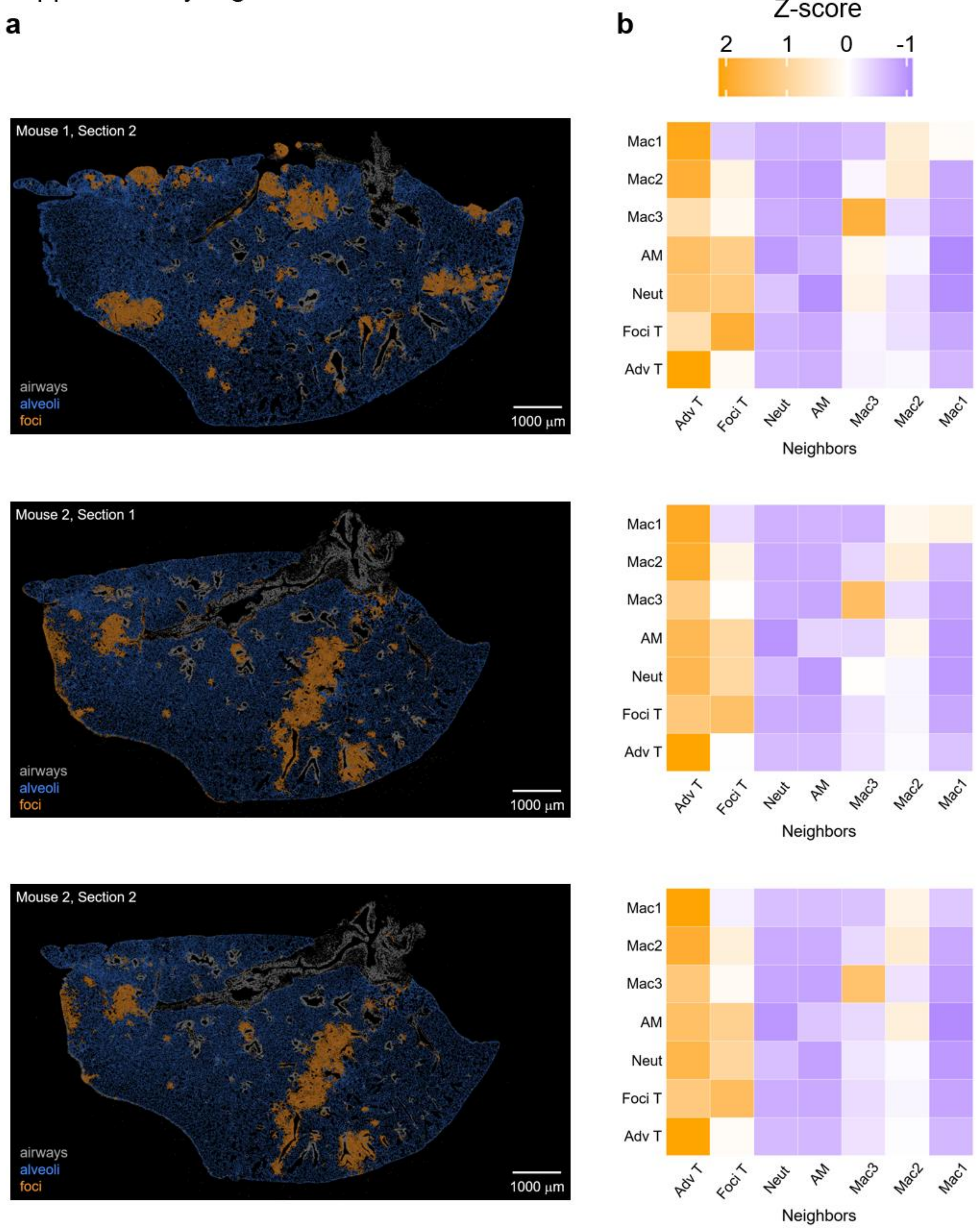


Supplementary Figure 7: Niche and cell neighborhood analysis of foci by section. (a) Images of 3 sections not shown in Figure 6, colored by niche. (b) Corresponding cell neighborhood heatmaps for foci displayed to right of each image and show that cells within foci in different sections have similar organization.

### Supplementary Figure 8

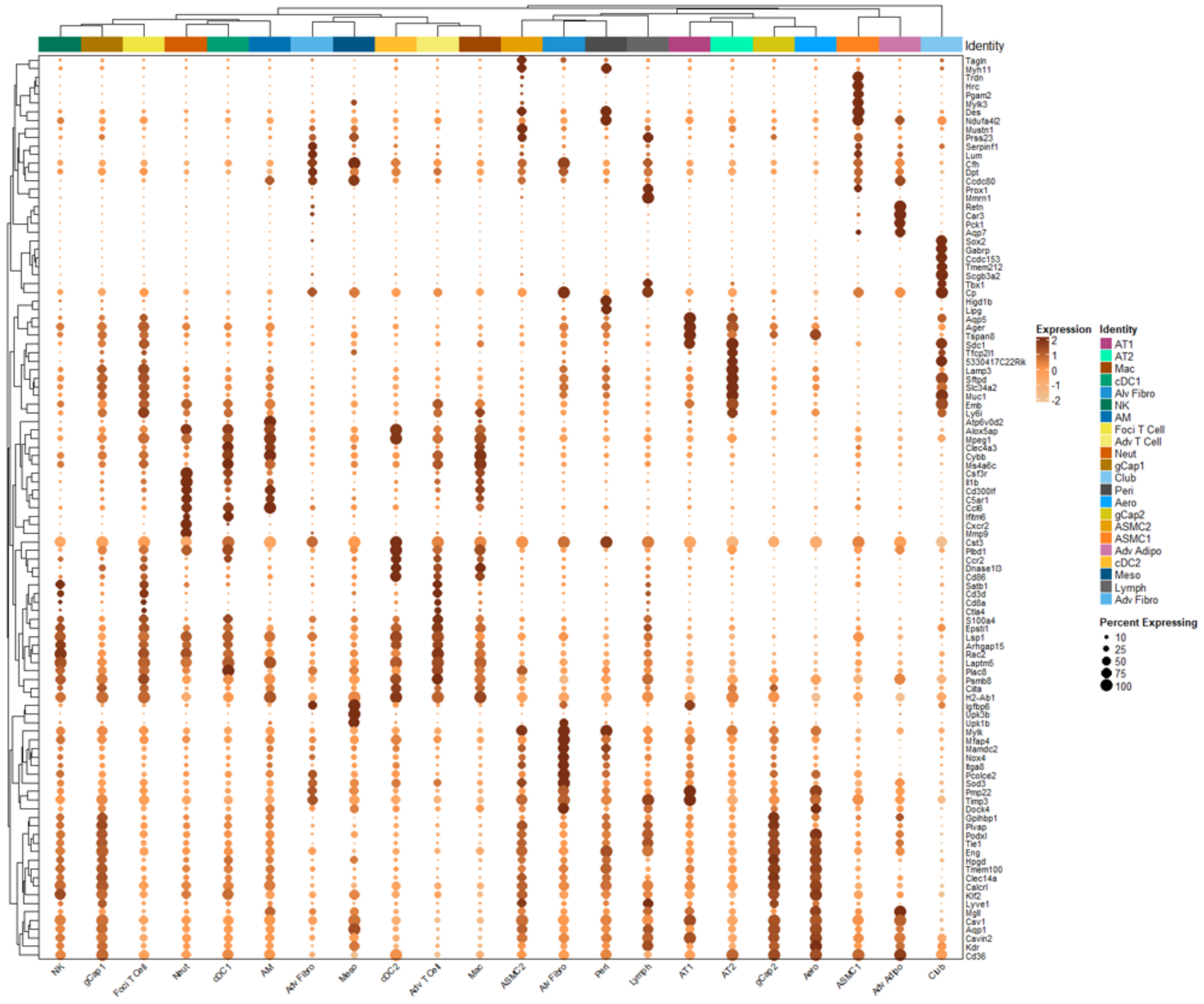


Supplementary Figure 8: Xenium panel genes used to annotate nearest-neighbor clusters. Dot plots show relative gene expression and frequency of expressing cells. Data are compiled from analysis of 2 sections each from 2 infected mice.

Supplementary Figure 9

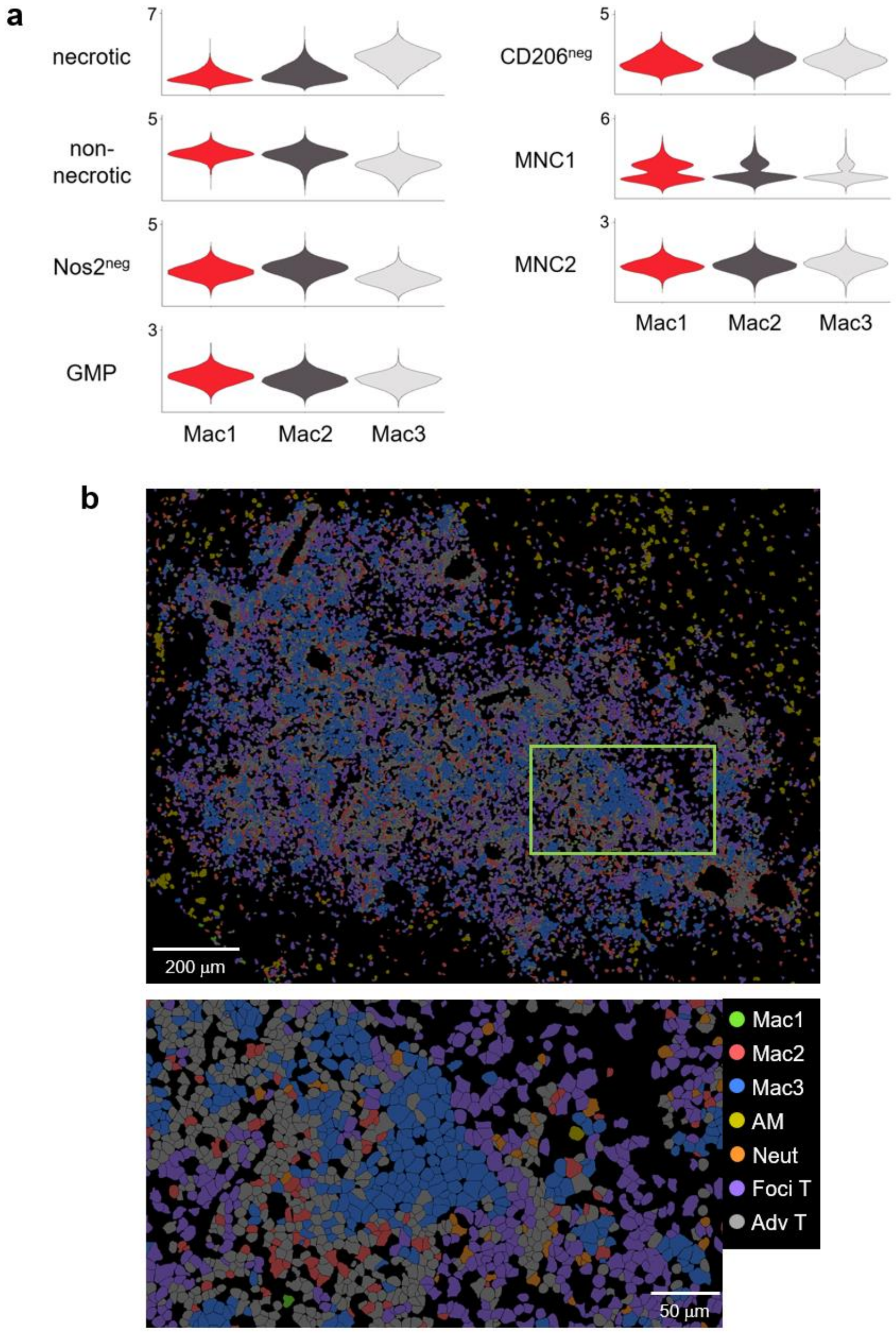


Supplementary Figure 9: Analysis of foci by spatial transcriptomics. (a) Violin plots of expression of the gene modules from Chai et al. ("necrotic" and "non-necrotic" granulomas), Pisu et al. ("Nos2$^{neg}$" interstitial macrophages), Trzebanski et al. (GMP-derived macrophages), Schyns et al. ("CD206$^{neg}$" interstitial macrophages), and Zheng et al. ("MNC1" and "MNC2"). Data are compiled from analysis of 2 sections each from 2 infected mice. (b) Higher magnification views of image in Figure 6 show all annotated cell populations in a representative focus of immune cells.
